# Whole-brain mapping reveals the divergent impact of ketamine on the dopamine system

**DOI:** 10.1101/2023.04.12.536506

**Authors:** Malika S. Datta, Yannan Chen, Shradha Chauhan, Jing Zhang, Estanislao Daniel De La Cruz, Cheng Gong, Raju Tomer

## Abstract

Ketamine is a multifunctional drug with clinical applications as an anesthetic, as a pain management medication and as a transformative fast-acting antidepressant. It is also abused as a recreational drug due to its dissociative property. Recent studies in rodents are revealing the neuronal mechanisms that mediate the complex actions of ketamine, however, its long-term impact due to prolonged exposure remains much less understood with profound scientific and clinical implications. Here, we develop and utilize a high-resolution whole-brain phenotyping approach to show that repeated ketamine administration leads to a dosage-dependent decrease of dopamine (DA) neurons in the behavior state-related midbrain regions and, conversely, an increase within the hypothalamus. Congruently, we show divergently altered innervations of prefrontal cortex, striatum, and sensory areas. Further, we present supporting data for the post-transcriptional regulation of ketamine-induced structural plasticity. Overall, through an unbiased whole-brain analysis, we reveal the divergent brain-wide impact of chronic ketamine exposure on the association and sensory pathways.

## INTRODUCTION

Ketamine is a schedule III (US Food and Drug Administration) substance with clinical applications as a dissociative anesthetic, as a pain management drug and, most recently, as a transformative fast-acting antidepressant^1–6^. Pharmacologically, ketamine is thought to act broadly in the brain^7^, most prominently as a non-competitive antagonist of the N-methyl-D-aspartate receptor (NMDAR)^8, 9^ but also as a blocker of hyperpolarization-activated cyclic nucleotide (HCN1)^10, 11^ channels and as a potential activator of opioid receptors^12, 13^. Recent studies in rodent models are unraveling the cellular and neural circuit underpinnings of ketamine’s complex action in the brain. For example, an antidepressant dose of ketamine was shown to promote spinogenesis and synaptogenesis in prefrontal cortical circuits to rescue the eliminated spines in a depression mouse model^14^. Another recent study showed that sub-hypnotic doses (50 and 100 mg/kg) of ketamine switches the spontaneous excitatory activity across the neocortex by suppressing the active neurons while activating the previously silent neurons, paralleling its dissociative property^15^. Acute ketamine administration also broadly impacts the dopaminergic modulatory system (via NMDAR antagonism^16, 17^), resulting in increased firing in the ventral tegmental area (VTA) dopamine (DA) neurons and enhanced DA release in the frontal cortex, striatum and nucleus accumbens^16–18^ (**Fig. 1a**).

**Figure 1.**
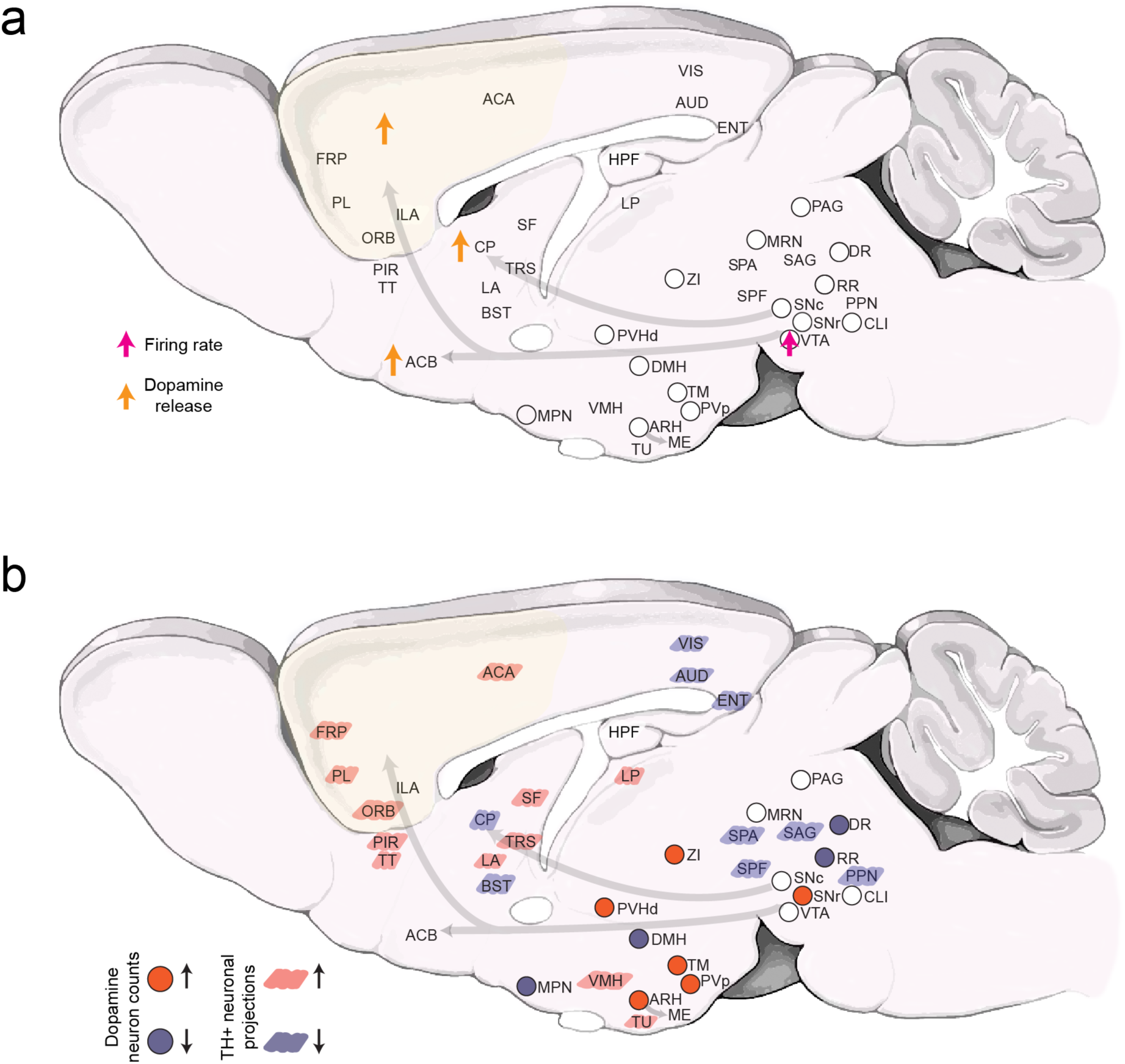
Brain-wide impact of ketamine exposure on the dopaminergic modulatory system. (a) Schematic summary of currently known alterations in dopaminergic neuronal activity and dopamine release after ketamine exposure^16, 17^. (b) Schematic summary (this study) of the brain-wide alterations in the DA system after 10 days of repeated ketamine (30 and 100 mg/kg) exposure. Up and down arrows indicate increase and decrease, respectively. Note that, for the 30 mg/kg ketamine treatment, statistically significant decreases in TH+ neuron counts were only observed in DR, DMH and MPN, and increase in ZI. The TH+ neuronal projections changes are only shown for the 100 mg/kg (10 days) treatment. Abbreviations used are standard Allen Brain Atlas terms, as follows. ACA: anterior cingulate area; ACB: nucleus accumbens; ARH: arcuate hypothalamic nucleus; AUD: auditory areas; BST: bed nuclei of the stria terminalis; CLI: central linear nucleus raphe; CP: caudoputamen; DMH: dorsomedial nucleus of the hypothalamus; DR: dorsal nucleus raphe; ENT: entorhinal area; FRP: frontal pole, cerebral cortex; HPF: hippocampal formation; IF: interfascicular nucleus raphe; ILA: infralimbic area; LA: lateral amygdalar nucleus; LP: lateral posterior nucleus of the thalamus; MPN: medial preoptic nucleus; MRN: midbrain reticular nucleus; ORB: orbital area; PAG: periaqueductal gray; PIR: piriform area; PL: prelimbic area; PPN: pedunculopontine nucleus; PVH: paraventricular hypothalamic nucleus; PVp: paraventricular hypothalamic nucleus, posterior part; RR: midbrain reticular nucleus, retrorubral area; SAG: nucleus sagulum; SF: septofimbrial nucleus; SNc: substantia nigra, compact part; SPA: subparafascicular area; SPF: subparafascicular nucleus; TM: tuberomammillary nucleus; TRS: triangular nucleus of septum; TT: taenia tecta; TU: tuberal nucleus; VIS: visual areas; VMH: ventromedial hypothalamic nucleus; VTA: ventral tegmental area; ZI: zona incerta.

In contrast, the long-term impact of chronic ketamine exposure on brain networks remains much less understood, with profound scientific and clinical implications^19, 20^. The antidepressant effect of ketamine is known to be transient, especially in treatment-resistant depression patients, thus often requiring maintenance treatments over years^19^. Additionally, the long-term recreational abuse has been associated with cognitive and sensory impairments^21–23^ and significant damages have been reported in the frontal, parietal, and occipital cortices in the brain^24^. Recent studies in mice have further revealed significant alterations in neocortical microcircuit synchrony after repeated exposure to ketamine^25, 26^. Therefore, with its broad clinical importance and increasing long-term abuse potential at higher doses, there is a considerable interest in understanding the molecular, cellular, and neural circuit alterations caused by long-term exposure to ketamine over a wide range of doses^19, 20^.

We sought to systematically investigate the brain-wide impact of chronic (*R,S*)-ketamine exposure on the entire dopaminergic system in mice. By utilizing a range of sub-hypnotic^15^ doses (30 and 100 mg/kg) and high-resolution whole-brain phenotyping of DA neurons, we show that chronic ketamine exposure over time results in divergent brain-wide changes in the DA neuron populations and their long-range projections to the prefrontal cortex and sensory areas. Further, we reveal the role of post-transcriptional regulation mechanisms in modulating the ketamine-induced structural plasticity in the DA system. Overall, through an unbiased whole-brain high-resolution mapping, we reveal the broad non-monotonic impact of ketamine on the brain-wide DA modulatory system.

## RESULTS

### High-resolution whole-brain phenotyping of ketamine-treated animals

We established a complete pipeline for whole-brain labeling, high-resolution imaging and comparative phenotyping of the entire dopaminergic modulatory system after 1, 5 and 10 days of daily (*R,S*)-ketamine (30 and 100 mg/kg) and saline control intraperitoneal (i.p.) injections (**Fig. 2**). The cellular toxicity of the ketamine exposure was assessed by α-activated caspase antibody staining after 10 days of 100 mg/kg daily i.p. injections, revealing no significant cell death in the brain (**Supplementary Fig. 1**). The locomotion of injected animals was video-recorded and quantified at 15’ and 60’ post injections (**Supplementary Fig. 2**). The 30 mg/kg group, but not 100 mg/kg, exhibited increasing (with days of exposure) locomotion sensitivity 15’-post injection.

**Figure 2.**
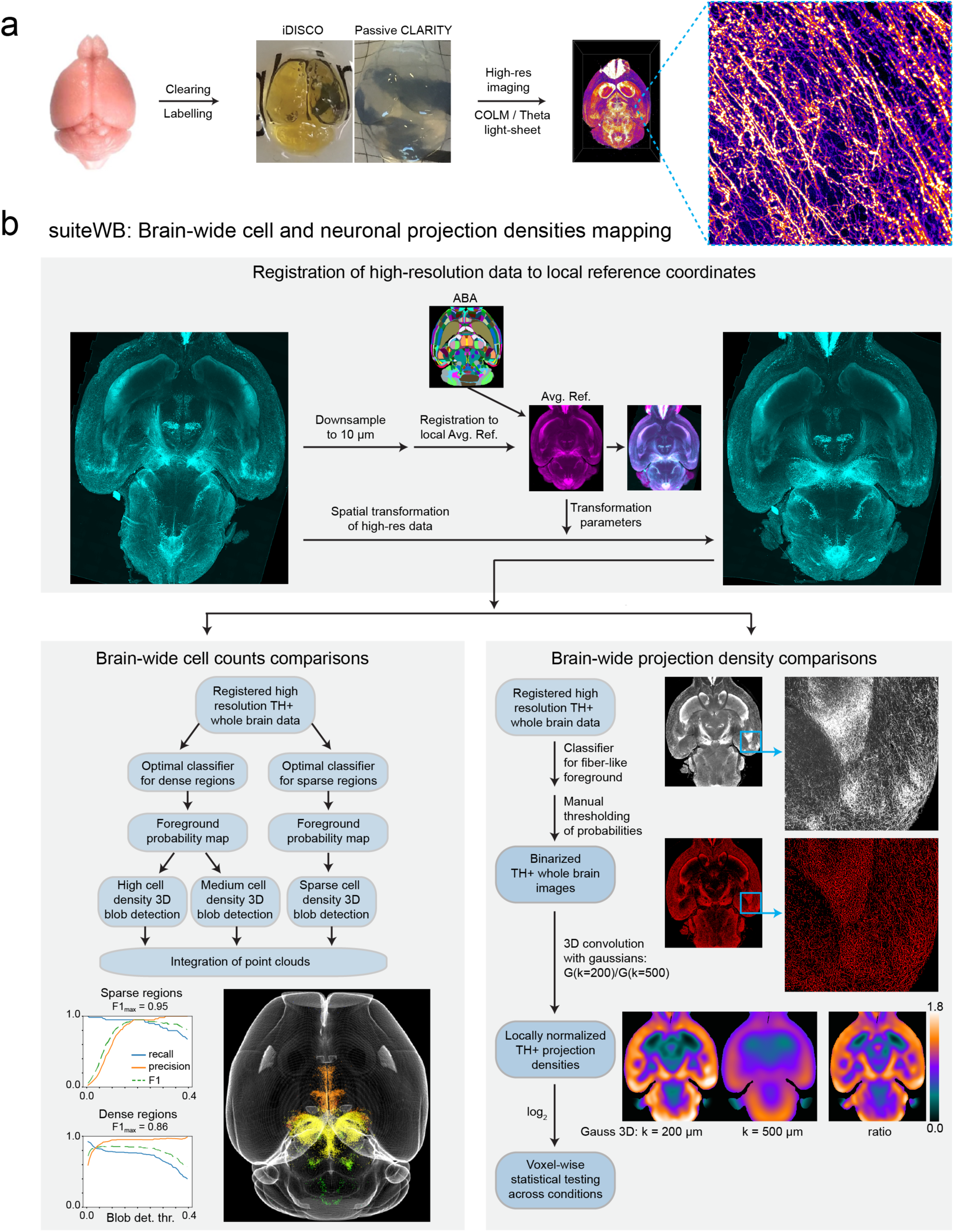
High-resolution whole-brain phenotyping of dopaminergic modulatory system. (a) Whole brain mapping of TH+ neurons by optimized labelling (α-TH antibody staining, TH mRNA staining and/or TH-CreER x tdTomato signal), clearing (iDISCO+ or passive CLARITY) and high-resolution imaging with COLM or LSTM light sheet microscopy. (b) A suite of tools (suiteWB) for high-resolution comparative phenotyping of neuron counts and their brain-wide projection, including registration of high-resolution whole-brain images to a local average reference brain (with mapped ABA reference annotations), accurate brain-wide multi-model cell segmentation approach and neuronal projections density estimation. F1 for sparse and dense region segmentations achieved are 0.95 and 0.86, respectively. Also see **Supplementary Videos 1-4** for volume renderings of the raw data and segmentation results.

The extracted intact brains were stained with α-tyrosine hydroxylase (TH; rate-limiting enzyme for DA synthesis), which is a widely used marker of DA neurons^27^ (**Fig. 2, Supplementary Video 1**). Note that even though DA may further get converted to other catecholamines (norepinephrine and/or epinephrine) in the downstream pathways, the distribution of these noradrenergic neurons is very well characterized and known to be localized within the hindbrain regions (pons and medulla)^28, 29^. Nevertheless, due to their brain-wide projections (e.g., to cerebral cortex, hippocampus, amygdala and hypothalamus), the precise identity of the TH+ neuronal projections (as DA or noradrenergic) may only be inferred as catecholaminergic^28, 29^.

For investigating the TH mRNA expression, we established and used a whole-brain staining method based on hybridization chain reaction^30, 31^, and also utilized a well-characterized inducible Cre line, TH-CreER^32^, crossed with a tdTomato reporter line (**Supplementary Fig. 3**). The intact brain samples were cleared with either iDISCO+^33^ or passive CLARITY^34^ methods, and imaged at high-resolution with COLM^34^ or light sheet theta microscopy (LSTM^35^) (**Fig. 2**).

Finally, we developed a set of accurate large data analysis methods (suiteWB, **Fig. 2b**) for high-resolution phenotyping of the entire dopaminergic system – both at the levels of TH+ cell bodies as well as TH+ brain-wide projections (**Fig. 2b**). To this end, we first generated a local average reference (from 7 brains), which was annotated by the registration of Allen brain atlas (ABA, ccfv3^36^) annotations. In addition, we developed a multi-model image segmentation approach to accurately detect the TH+ cell bodies and their brain-wide projections (**Fig. 2b, Supplementary Video 3**). ANOVA and two-sided Mann Whitney U tests with Bonferroni correction were utilized for statistical comparisons. Overall, suiteWB methods allow accurate high-resolution phenotyping of the brain-wide structural plasticity.

### Dosage-dependent divergent impact of chronic ketamine exposure on the DA domains

We generated high-resolution whole-brain maps of TH+ neurons after 1, 5, and 10 days of daily ketamine and saline i.p. injections (**Fig. 3, Supplementary Video 3**). Using suiteWB, TH+ neuron counts were calculated across all the brain regions (**Fig. 3a**) and were statistically compared across treatment groups at multiple scales by utilizing different graph-cut levels of the hierarchical ABA annotation tree. Robust, statistically significant alterations were only detected after 10 days of ketamine exposure for both 30 and 100 mg/kg ketamine treatment groups, therefore, 1- and 5-days treatment datasets were not analyzed further. In addition, as expected, 100 mg/kg treatment group exhibited much more changes than the 30 mg/kg treatment group (**Fig. 3**).

**Figure 3.**
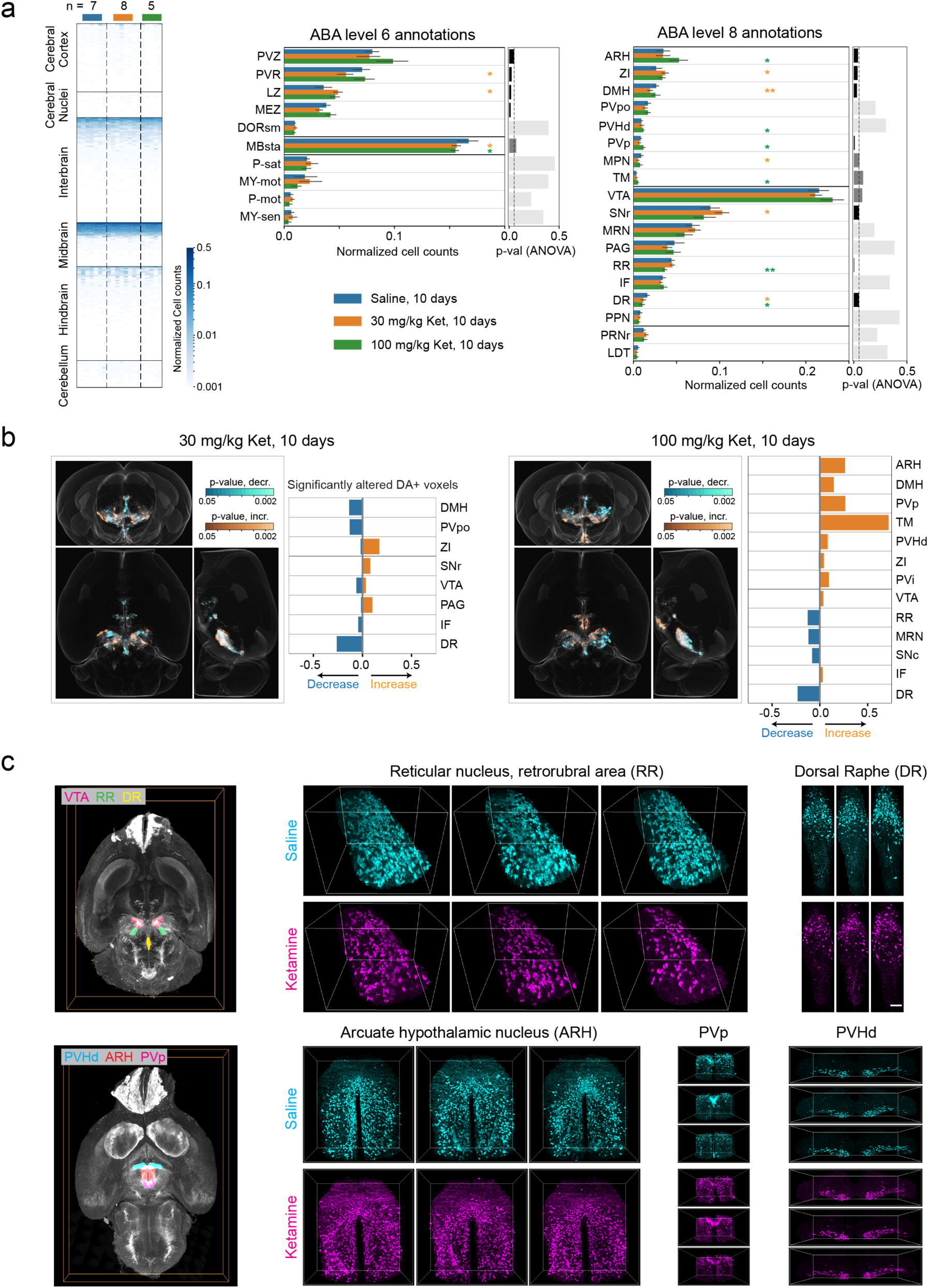
Repeated ketamine exposure results in divergent brain-wide changes in the dopaminergic modulatory system. (a) Brain-wide heatmap of the TH+ cell counts in ABA ROIs after 10 days ketamine (30 mg/kg and 100 mg/kg) and saline (Sal) i.p. injections. Independent biological replicates of n=7 for Sal, 8 for 30 mg/kg ketamine and 5 for 100 mg/kg ketamine treated groups. The regional TH+ neuron counts were normalized by respective total number of TH+ neurons in brain. Orange, green, and blue bars quantify TH+ cells in 30 mg/kg ketamine, 100 mg/kg ketamine and saline-treated brains, respectively. Two-sided Mann-Whitney U tests with Bonferroni Correction were performed to calculate the statistical significance. * and ** indicate p-values < 0.05 and < 0.01, respectively. ANOVA tests p-values are plotted as bar plots on the right. (b) Whole brain orthogonal projections visualizing voxel-by-voxel differences in TH+ neuron densities after 10 days of 30mg/kg ketamine (n=8) and 100mg/kg ketamine (n=5), compared with saline (n=7) controls. Cyan and Orange visualize p-values (two-sided Mann-Whitney U test) for decreases and increases, respectively. Note that the decreases and increases in the densities are identified by comparing the sample averages of the treated and control groups. The bar plots quantify the fraction of DA neurons containing voxels (within an ABA ROI) that show significant alterations. (c) Representative volume renderings from 3 different brain samples, each for ketamine (100 mg/kg, 10 days) and saline samples, showing decreases in RR (Reticular nucleus, retrorubral area) and DR (Dorsal raphe), and increases in ARH (Arcuate hypothalamic nucleus), PVp (Periventricular hypothalamic nucleus, posterior part) and PVHd (Periventricular hypothalamic nucleus, descending division). Renderings on the left visualize the spatial locations of these regions in the brain.

At a higher-level annotation (i.e., larger brain regions ROIs; level 6), we found an overall dose-dependent statistically significant decrease in TH+ neuron counts within the behavior-state related mid-brain regions (MBsta) and, conversely, an overall increase within the hypothalamic lateral zone (LZ) in both the 30 and 100 mg/kg (10 days) treatment groups (**Fig. 3a**). In addition, for the 100 mg/kg group, we observed a tendency for increase within the hypothalamic periventricular zone (PVZ) and, for the 30 mg/kg group, significant decrease within the periventricular region (PVR) (**Fig. 3a**). Next, we compared the brains at lower-level (level 8) graph cut of the ABA annotation tree. For both 30 and 100 mg/kg treatment groups, we observed a robust decrease in the dorsal raphe (DR^37^) and increase in the lateral hypothalamic region zona incerta (ZI^38^). In addition, in the 100 mg/kg treatment group, we observed a robust decrease in the reticular nucleus retrorubral area (RR^39^) and increases in the arcuate hypothalamic nucleus (ARH^40^), the periventricular hypothalamic nucleus posterior part (PVp), tuberomammillary nucleus (TM) and the periventricular hypothalamic nucleus descending division (PVHd) (**Fig. 3a**). Whereas, in the 30 mg/kg treatment group, we found significant increases in mid-brain region SNr (Substantia nigra reticular part) and a significant decrease in the dorsomedial nucleus of the hypothalamus (DMH) and the medial hypothalamic region MPN^41^ (preoptic nucleus). These brain-wide alterations were further validated by an independent voxels-based (20x20x20 µm^3^ sampling) brain parcellation approach (**Fig. 3b**). Representative example volume renderings are shown for RR, DR, ARH, PVp and PVHd (**Fig. 3c**, 100 mg/kg ketamine-treated group).

Overall, an unbiased whole-brain comparison of TH+ neuron counts revealed divergent and dosage-dependent brain-wide impact of chronic ketamine exposure (**Fig. 1**) - increases within multiple hypothalamic domains (e.g., ARH, containing TH+ DA neurons with orexigenic function in energy homeostasis^40^) and decreases within the behavioral state related midbrain regions (e.g., DR, containing TH+ DA neuron which modulate the social isolation behaviors^37^; RR, containing TH+ DA neurons with role in fear and aversive signaling^39^).

### Untranslated TH mRNA+ neurons facilitate ketamine-induced cellular plasticity

We sought to further investigate the mechanistic basis of the chronic ketamine-induced brain-wide cellular plasticity. Utilizing our whole-brain mRNA labeling protocol, we first mapped the expression of TH mRNA (**Fig. 4a, Supplementary Video 4**), revealing a much broader distribution than the corresponding TH protein. Such discrepancy in the TH mRNA/protein expression (**Supplementary Video 5**) suggests a potential role for post-transcriptional mechanisms in rapid modulation of the brain-wide DA system. To test this hypothesis further, we utilized a well-characterized inducible TH-CreER transgenic line^32^ (**Supplementary Fig. 2**), crossed with a tdTomato reporter line, to investigate if the newly acquired TH+ neurons originated from the untranslated TH mRNA+ neurons. TH-CreER induction (by 4-OHT i.p. injections) was performed 1-week prior to the start of the ketamine exposure, thus, permanently labeling the pre-treatment TH mRNA+ neurons by tdTomato. After 10 days of chronic ketamine (100 mg/kg) exposure, the brains were harvested, cleared with passive CLARITY^34^, and labeled with α-TH antibody for a direct comparison of the after-treatment TH+ neurons with before-treatment TH mRNA+ neurons, within the exact same brain samples. As shown in **Fig. 4b**, we found a significant increase in TH protein+/tdTomato+ co-labeled neurons (α-TH ∩ tdTomato) in the ketamine-treated group (compared to saline controls), while the overall number of tdTomato+ neurons remained unchanged, consistent with unchanged TH mRNA+ neurons (**Fig. 4b**). Conversely, we observed a decrease in TH protein+/tdTomato+ co-labeled neurons in the midbrain regions (which showed reduction in TH+ DA neuron counts), while the tdTomato+ and TH mRNA+ neuron counts remain unchanged compared to the saline controls (**Fig. 4c**). Altogether, these results suggest that the cellular plasticity in the dopaminergic system may be facilitated by the existence of a much larger pools of untranslated TH mRNA+ neurons to rapidly modulate the number of available TH+ DA neurons in various regions of the brain.

**Figure 4.**
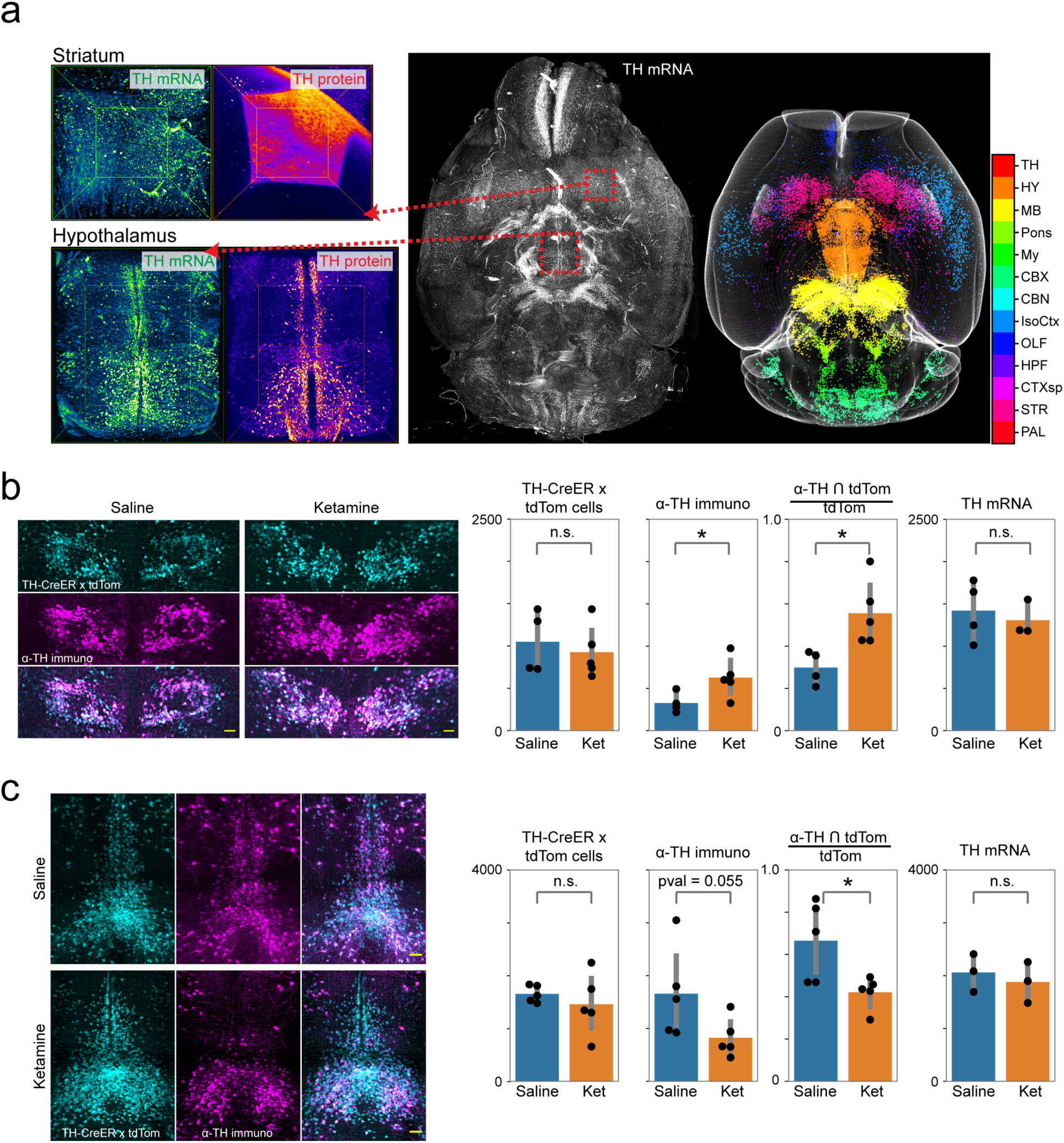
Untranslated TH mRNA+ neurons facilitate adaptations in the dopamine system. (a) Whole brain mapping of TH mRNA, showing much broader expression than the TH protein. Color code corresponds to different ABA ROIs, as listed in adjoining color bar. (b-c) Co-labeling of TH-CreER x tdTomato signal induced with 4-OHT injections one week before the start of ketamine i.p. injections and α-TH immunostaining signal capturing TH protein expression post-treatment (10 days). Left to right, bar plots compare the number of tdTomato+ neurons, α-TH immunostained neurons, their intersection and the number of TH mRNA+ neurons. (b) and (c) show field-of-views that include paraventricular hypothalamic nucleus and dorsal raphe regions, respectively. Two-sided Mann-Whitney U tests were performed for statistical significance calculations. * indicates <0.05 p-value. All scale bars are 100 µm. Also see **Supplementary Videos 4** and **5** for whole brain volumetric renderings.

### Altered long-range TH+ projections after chronic ketamine exposure

Taking advantage of the high-resolution of our datasets, we sought to map the brain-wide changes in TH+ neuronal projections. Note that TH+ neuronal projections may be DA or noradrenergic, therefore they can only be inferred as catecholaminergic^28, 29^. Using the suiteWB pipeline, we estimated the projection densities in the ketamine (100 mg/kg, 10 days) treated and saline control whole-brain datasets and performed a voxel-by-voxel (at 25x25x25 µm^3^ sampling) statistical comparisons across groups. As shown in **Fig. 5, Supplementary Video 6**, chronic ketamine exposure resulted in robust brain-wide changes in TH+ projections. We observed increased densities in multiple associative cortical regions including the prefrontal cortex (PFC)-related prelimbic area (PL, **Supplementary Video 7,** **Fig. 5c**), orbital area (ORB, **Supplementary Video 8,** **Fig. 5c**), frontal pole (FRP, **Fig. 5a**) and anterior cingulate area (ACA, **Fig. 5a**), and posterior parietal association area (PTLp, **Fig. 5a**). Further, the lateral amygdala (LA, **Supplementary Video 9,** **Fig. 5c**), which is crucial for processing of threatening stimuli and fear behavior, specific regions of the lateral septal complex (septofimbrial nucleus (SF), involved in reward and reinforcement) and the triangular nucleus of septum (TRS) also showed increased projection densities (**Fig. 5a**). Furthermore, brain regions receiving olfactory inputs, including the taenia tecta (TT) and piriform area (PIR), and regions of the thalamus (lateral posterior nucleus of the thalamus (LP)) and hypothalamus (VMH and TU) also showed increases in the TH+ neuronal innervations.

**Figure 5.**
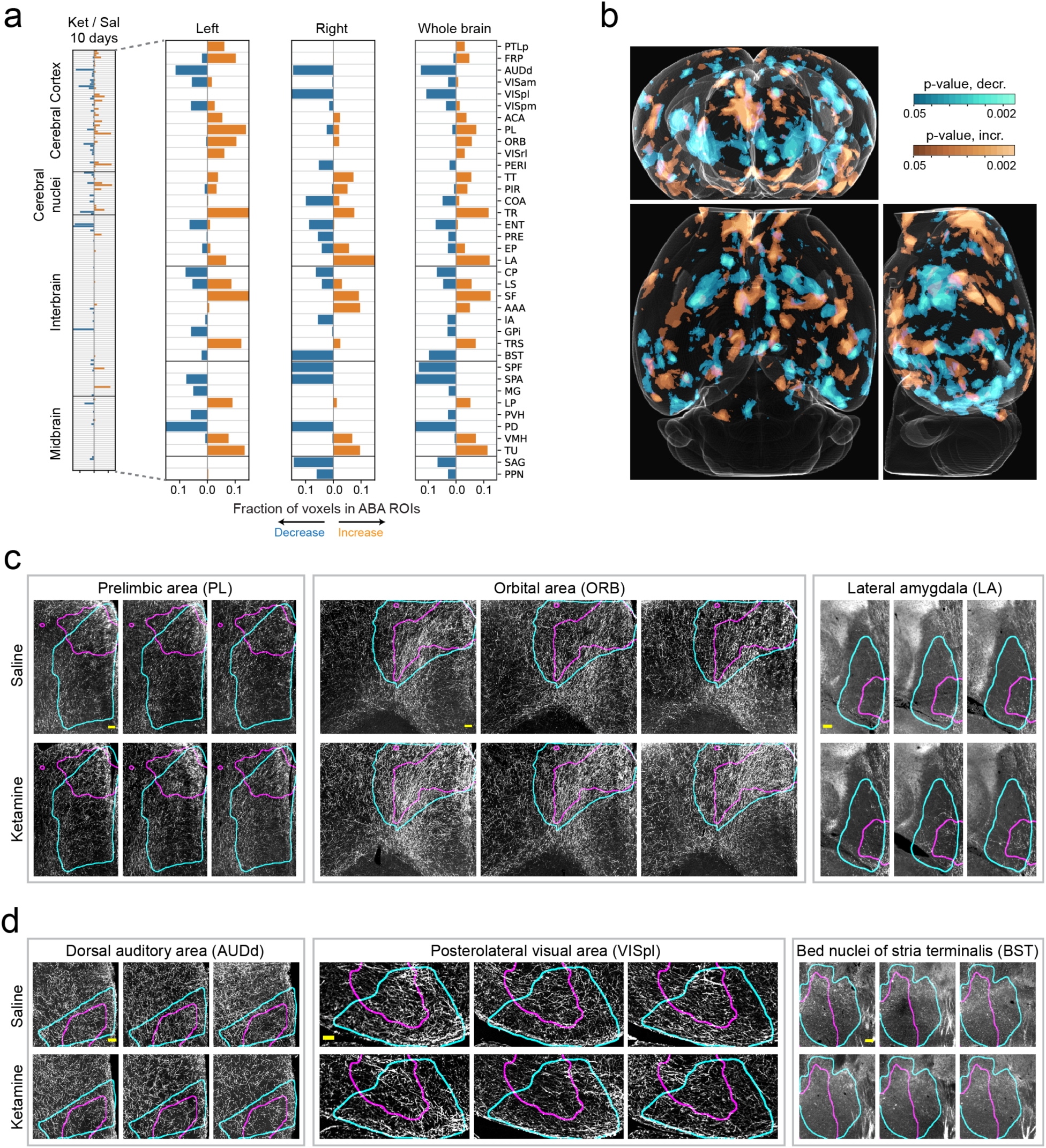
Brain-wide changes in TH+ neuronal projections after chronic ketamine exposure. (a) Stacked barplots of fractions of ABA ROI voxels with significantly increased (orange) and decreased (blue) TH+ neuronal projection densities after 10 days of ketamine exposure. Independent biological replicates as follows: n (Ket, 10d) = 5, n (Sal, 10d) = 7. (b) Whole brain orthogonal projections visualizing voxel-by-voxel differences (independent of ABA) in TH+ neuronal projection densities. Orange and Cyan represent significant increases and decreases, respectively. (c-d) Representative examples of ABA ROIs showing increases (c) and decreases (d) in TH+ neuronal projections. Cyan and Magenta curves outline the ABA ROIs and areas with significantly changed projection densities. Images shown are representative single optical planes. Also see **Supplementary Video 6** for rendering of heatmaps, **Supplementary Videos 7-12** for complete volumes. All scale bars are 100 µm.

In contrast, the dorsal auditory area (AUDd, **Fig. 5d**, **Supplementary Video 10**), postero-lateral visual area (VISpl, **Fig. 5d**, **Supplementary Video 11**), entorhinal area (ENT) and presubiculum (PRE, involved in spatial information processing; **Fig. 5a**) exhibited decreased TH+ projection densities. Striatum sub-regions caudoputamen (Cp) and bed nuclei of the stria terminalis (BST/BNST, **Fig. 5d**, **Supplementary Video 12**) showed reductions consistent with the loss of TH+ neurons in DR, RR and SNc (**Fig. 3**). Finally, thalamic regions subparafascicular area (SPA) and subparafascicular nucleus (SPF), and midbrain regions nucleus sagulum (SAG) and pedunculopontine nucleus (PPN) also showed decreased projections.

Overall, consistent with the divergent changes in TH+ neuron counts in the midbrain and hypothalamic dopaminergic domains (**Fig. 3**), chronic ketamine exposure resulted in increased TH+ neuronal projection densities in the associative brain centers, including PFC-related regions, and decreased innervations of visual, auditory, and spatial information processing regions (**Fig. 5**).

## DISCUSSION

Utilizing an unbiased, high-resolution whole-brain mapping approach, we revealed hitherto unknown divergence in the adaptability of the DA system to repeated ketamine exposure (summarized in **Fig. 1b**) – reduced number of TH+ neurons within the behavior state-related mid-brain domains and increased within the hypothalamic domains, along with altered long-range innervation of the association and sensory areas by TH+ neuronal projections. Such structural plasticity of brain-wide modulatory system may facilitate significant reconfiguration of the neuronal networks in response to external stresses such as ketamine exposure or diseased conditions (e.g., schizophrenia), to eventually result in long-lasting cognitive behavioral changes. Note that, DA may further get processed in the noradrenergic neurons, but their cellular distribution is well known to be restricted to specific domains in hindbrain regions^28, 29^. However, the TH+ neuronal projections may still not be precisely identified as DA or noradrenergic by TH immunolabeling^28, 29^. In addition, the loss of TH protein in DA neurons may not result in a permanent loss of DA neurons, nevertheless, it does indicate the loss of DA producing capacity due to the required role of TH in the L-DOPA (DA precursor) biosynthesis.

We also show that the adaptability of the DA system is facilitated by a large pool of neurons that stably maintain translationally suppressed TH mRNA (**Fig. 4**). These findings were validated by a direct comparison of the newly recruited or lost TH+ DA neurons with the before-exposure TH mRNA expressing neuron populations by utilizing an inducible TH-CreER transgenic mice line (**Fig. 4**). Mechanistically, these observed changes in DA system may, in part, be the result of repeated ketamine-induced activation/deactivation (indirect, via NMDAR antagonism) of the DA neurons due to the heterogeneous neuronal inputs. Intriguingly, the ketamine-induced cellular plasticity is distinct from the previously reported neurotransmitter phenotypic plasticity of the hypothalamic DA neurons after changed day/night durations in rodents, and included loss of TH mRNA expression^42^. Overall, our data suggest that the maintenance of translationally suppressed mRNA, even though energetically costly process, may allow for faster brain-wide adaptations to various external stresses.

The development and use of unbiased, high-resolution whole-brain phenotyping of the entire DA system allowed discovery of the divergent impact of chronic ketamine exposure. Such non-monolithic brain-wide impact further underscores the need for unbiased investigations of on/off-target effects of ketamine treatment at a range of doses, as well as the urgency to develop targeted pharmacological intervention approaches (e.g., focused ultra sound-based approaches^43, 44^) for treatment of complex brain disorders.

## Author Contributions

M.S.D., Y.C. and R.T. conceptualized and designed the project. M.S.D. performed most of the experiments. J.Z. performed mRNA staining experiments. E.D.D. contributed with animal husbandry and general experimental support. S.C. contributed to all the imaging experiments, and C.G. contributed to ClearScope imaging. Y.C. and R.T. developed the data analysis framework and code, and with M.S.D. analyzed the data. R.T., M.S.D. and Y.C. wrote the paper with inputs from all authors. R.T. supervised the project.

## Data availability statement

All datasets will be made available on request.

## Code availability

All code will be made available publicly as a GitHub repository.

## Methods

### Animals

Male TH-2a-CreER^32^ mice were acquired from Dr. David Ginty’s lab and were bred with B6;129S6-*Gt(ROSA)26Sor^tm14(CAG–tdTomato)Hze^*/J (Ai14; JAX Strain #:007908). All mice were group-housed in a 12:12 light:dark cycle at 22°C. Food and water were provided *ad libitum. (R,S)-* ketamine i.p. injections were performed during the light phase. All experimental procedures were approved by the IACUC at Columbia University. Locomotor activity was recorded after 1, 5, and 10 days of treatment. The mice were placed within a novel home cage and were recorded (camera: GoHZQ, 1920 x 1080 pixels,30 fps transmission rate) 15 minutes and 1 hour post-intraperitoneal ketamine injections. The total distance traveled was quantified using the ANY-maze tracking software (ANY-maze, RRID:SCR_014289, Stoelting, Wood Dale, IL, United States).

### Drugs

4-hydroxytamoxifen (4-OHT; Sigma, H7904) was dissolved in corn oil/ethanol (90% corn oil, 10% ethanol) via the use of a vortex, ultrasonication, and 55°C heating for <15 min. Mice were intraperitoneally injected with 2 mg of 4-OHT. *(R,S)*-ketamine (Covetrus) was used for all ketamine exposure experiments. 3 and 10 mg/mL stocks were prepared in saline (0.9% NaCl)^45^. One dose of 30 mg/kg or 100 mg/kg of *(R,S)*-ketamine was intraperitoneally injected in a 24 hour period for 1 day, 5 days, or 10 days treated animals. 8-10 weeks old mice were used for all experiments.

### Whole brain clearing and labeling

For iDISCO clearing, the iDISCO+ protocol was followed as previously described^33^. The brains were pre-treated with methanol, placed in 66% dichloromethane/ 33% methanol overnight, bleached with 5% H_2_O_2_/methanol, and then rehydrated. Afterwards, the whole brains were permeabilized for 2 days at 37°C in 1x phosphate buffered saline (PBS), 0.2% Triton X-100, 0.3 M glycine, and 20% dimethyl sulfoxide (DMSO). The brains were blocked with 1xPBS, 0.2% Triton X-100, 10% DMSO, 6% donkey serum for 2 days at 37°C, followed by incubation in 1xPBS, 0.1% Triton X-100, 3% donkey serum, and a 1:200-1:500 dilution of the primary antibody sheep α-TH (ab113, Abcam) for 10 days. After washing in 1xPBS/0.1% Triton X-100, the brains were placed in the secondary antibody solution containing 1xPBS/0.1% Triton X-100/3% donkey serum and a 1:1000 dilution of donkey anti-sheep 647 (A-21448, Thermofisher) for 10 days.

We used the passive CLARITY method as detailed previously^34^. The tissue was first encapsulated in a hydrogel monomer (HM) solution consisting of 1% (wt/vol) acrylamide, 0.05% (wt/vol) bisacrylamide, 4% paraformaldehyde, 1x phosphate buffered saline (1xPBS), deionized water, and 0.25% of thermal initiator (VA-044, Fisher Scientific), followed by clearing with SBC buffer (4% (wt/vol) SDS, 0.2 M boric acid, pH 8.5 and deionized water) at 37°C with shaking. The SBC buffer was replaced every 2 days. After clearing, the SBC buffer was washed off with 0.2 M boric acid pH 8.5 with 0.1% Triton X-100. The cleared tissue was immunostained in 0.2 M boric acid pH 7.5 with 0.1% Triton X-100. The final refractive index matching was performed in RapiClear (SunJin lab, RI=1.47).

For HCR-FISH, we used split-initiator DNA probes (IDT) for detecting the tyrosine hydroxylase (Th, NM_012740.3) mRNA. During probe hybridization/detection, the brains underwent 1) equilibration with 5xSSCTw buffer, 2) acetylation with 0.25% acetic anhydride solution, 3) equilibration with probe hybridization buffer (30% formamide, 5xSSC, 0.5 mg/mL yeast tRNA, 10% dextran sulfate), and 4) probe incubation (50 nM) in probe hybridization buffer at 37C. The time/amount of solution varied depending on the thickness of the brain slices. The slices were then washed in probe wash buffer (30% formamide, 5X SSC, 9 mM citric acid, and 0.1% tween 20) at 37°C as well as two rounds of washes in 5xSSCTw. Afterwards, the slices were equilibrated in amplification buffer (5xSSC, 10% dextran sulfate, and 0.1% Tween-20), and then incubated in amplification buffer with 50-150 nM of hairpins H1 and H2 conjugated with AF-647 (Molecular Instruments). The hairpin/amplification buffer mixture was then washed off with several rounds of 5xSSCTw.

For activated caspase staining, the TH-2a-CreER;Ai14 brains were sliced with a vibratome (Leica VT1000 S Vibrating blade microtome) into 50 µm sections. These sections were placed in a blocking buffer (1xPBS/0.1% bovine serum albumin/0.1% Triton X-100) for 30 minutes and then incubated in a 1:200-1:400 dilution of the primary antibody, rabbit anti-active caspase-3 (BD Biosciences, catalog no. 559565), with 1xPBS/0.5% bovine serum albumin /0.1% Triton X-100 overnight. After washing in 1xPBS/0.1% Triton X-100, the sections were incubated, for two hours, in a 1:500 dilution of the secondary antibody, goat anti-rabbit 647 (ThermoFisher, Catalog #A-21245) with 1xPBS/0.5% bovine serum albumin /0.1% Triton X-100. The secondary antibody was washed off in 1xPBS/0.1% Triton X-100, and the slices were incubated in a 1:1000 dilution of a 1 mg/mL stock of DAPI with 1xPBS for 15 minutes. After washing off with 1xPBS/0.1% Triton X-100, the slices were mounted in 65% glycerol.

### Imaging

All imaging experiments were performed with CLARITY-optimized light sheet microscopy (COLM)^34^ or ClearScope (MBF Biosciences)^35^ Olympus 10x/0.6NA/8 mmWD or ASI 16x/12mmWD detection objectives were used for most of the whole brain imaging with COLM. Olympus Macro 4×/0.28NA or Nikon 20x/1.0NA were used for ClearScope imaging.

### suiteWB: Image Registration

We first developed a fast and efficient multistep multiresolution 3D image registration tool by using Mutual Information^46^ as the similarity metric. The registration starts by rigid transformation (with 6 degrees of freedom for rotation and translation) of the moving image (i.e. the image being aligned) to roughly align with the reference image. Next, the moving image is affine transformed (12 degrees of freedom) to account for shearing and shrinking artifacts introduced by labeling, tissue clearing and imaging. Finally, we used a uniform grid of control points and third-order B-splines are used for non-rigid local transformations. Affine and nonrigid steps were done at three different resolutions. The algorithms were implemented with ITKv4^47^ registration framework. All α-TH stained whole brain images were registered at 10 µm resolution to a local average reference brain (generated from 7 α-TH whole brain images), and the resulting spatial transformation parameters were applied to high-resolution datasets. ABA annotations were registered one time to the local average reference brain.

### suiteWB: Multi-model segmentation pipeline

Segmentation of whole brain images presents unique challenges of high spatial variations in signal-noise ratios (SNR), object densities and varied image artifacts. Multiple open-source tools exist for cell segmentation^48–50^, however, some of them are optimized for confocal images with smaller data sizes, others utilize deep learning models with pre-trained parameters that do not accurately generalize to images with different signal quality distribution and also require dense annotation training datasets. We chose to develop a semi-supervised multi-model learning approach to address the challenges of whole brain segmentation. This approach does not require dense annotated training and also allows the flexibility of using different optimal parameter sets for different regions (i.e. with dense or sparse object densities). We utilized open-source toolkit ilastik^51^ for semi-supervised learning of classifiers based on image features.

The high-resolution registered whole brain images were split into densely and sparsely populated regions by visual inspection, followed by pixel classification to generate probability maps. For the sparse region, a standard pixel classification workflow in ilastik was applied to generate cell probability maps. First, image features were extracted by using multiple types of filters with different kernel sizes, including intensity filters (Gaussian Smoothing), edge detection filters (Laplacian of Gaussian, Gaussian Gradient Magnitude, Difference of Gaussians), and texture detection filters (Structure Tensor Eigenvalues, Hessian of Gaussian Eigenvalues). The image features were then used to estimate the probabilities of pixels belonging to a cell or background. For denser regions, we utilized the Autocontext workflow in ilastik. In the first step, the image pixels were classified into five categories: empty space, brain background, fibers (neuronal projections), cell cytoplasm and cell nucleus. In the second step, the classification results were combined with image features (with filters described above for sparse regions) to generate probability maps of cells against background. Next, we applied blob detection on the dense and sparse probability maps separately to detect cells. This was achieved by using the difference of Gaussian (DOG) algorithm (Python scikit-image package^52^. To optimize memory usage, images were processed block-wise with carefully resolved boundary conditions. Cells detected in dense and sparse regions were merged and were further tuned by using Napari viewer. F1 for sparse and dense region segmentations reached 0.95 and 0.86, with precision 0.889 and 0.976 respectively, compared to human-annotated datasets (**Fig. 2**). For comparison, ClearMap^48^, applied to immediate early gene datasets, reported a precision of 0.83 and 0.75 compared to two human annotators.

For colocalization analysis of the multi-channel (i.e. tdTomato and α-TH immunostaining signal) CLARITY images, cell detection (as described above) was applied separately to different channels. The full cell body labels were generated by taking the union of the detected cells in the two channels that were less than 25µm apart. The cell identities (i.e. if expressing tdTomato or α-TH or both) were assigned by manually validated thresholding of probability maps of each channel.

Neuronal projection segmentation was performed by semi-supervised pixel classification by using image features (as discussed above) to generate the probability maps distinguishing projections and background. The probability threshold for binarization was determined by careful inspection throughout various regions of the brain. The binarized data was then convolved with 3D Gaussian kernels of 200 µm and 500 µm. The ratio of the two gaussians resulted in locally normalized neuronal projection densities.

Statistical comparisons between Ketamine and Saline treated samples were done with two-sided Mann Whitney U test (using Python package Scipy^53^) on normalized cell counts (cell count in a region divided by total cell count of that whole brain) and cell densities (estimated by 3D Gaussian kernel of size 50 µm in each dimension).

## Acknowledgements

We are grateful to Jordan Hamm for critically reading the manuscript, and to Christine Denny, Rae Silver and Darcy Kelley for advice throughout the project. We thank Vivek Kumar for advice on the HCR probe design and general discussions. We are grateful to Briana Chen and Christine Denny for helping with the ANY-maze software. This work was supported by NIH grant DP2MH119423 and Columbia University Arts & Sciences startup grant to R.T.

