## Supplementary Figures and Video legends for "Whole-brain mapping reveals the divergent impact of ketamine on the dopamine system"

**Table of contents**

**Supplementary Fig. 1 | Activated caspase immunostaining.**

**Supplementary Fig. 2 | Locomotion behavior quantification.**

**Supplementary Fig. 3 | Characterization of TH-CreER transgenic line.**

**Supplementary Video 1 | High-resolution whole brain TH immunostaining and imaging.**

**Supplementary Video 2 | Whole brain image registration.**

**Supplementary Video 3 | Volumetric rendering of whole brain TH+ cell segmentation.**

**Supplementary Video 4 | Volumetric rendering of whole brain TH mRNA+ cell segmentation.**

**Supplementary Video 5 | Comparison of whole brain TH mRNA+ (left) and TH protein+ neuronal cell bodies.**

**Supplementary Video 6 | Whole brain heatmaps of alterations in TH+ neuron counts and projections.**

**Supplementary Video 7 | TH+ projection density changes in the prelimbic area.**

**Supplementary Video 8 | TH+ projection density changes in the orbital area.**

**Supplementary Video 9 | TH+ projection density changes in the lateral amygdala.**

**Supplementary Video 10 | TH+ projection density changes in the dorsal auditory area.**

**Supplementary Video 11 | TH+ projection density changes in the postero-lateral visual area.**

**Supplementary Video 12 | TH+ projection density changes in the Bed nuclei of the stria terminalis.**

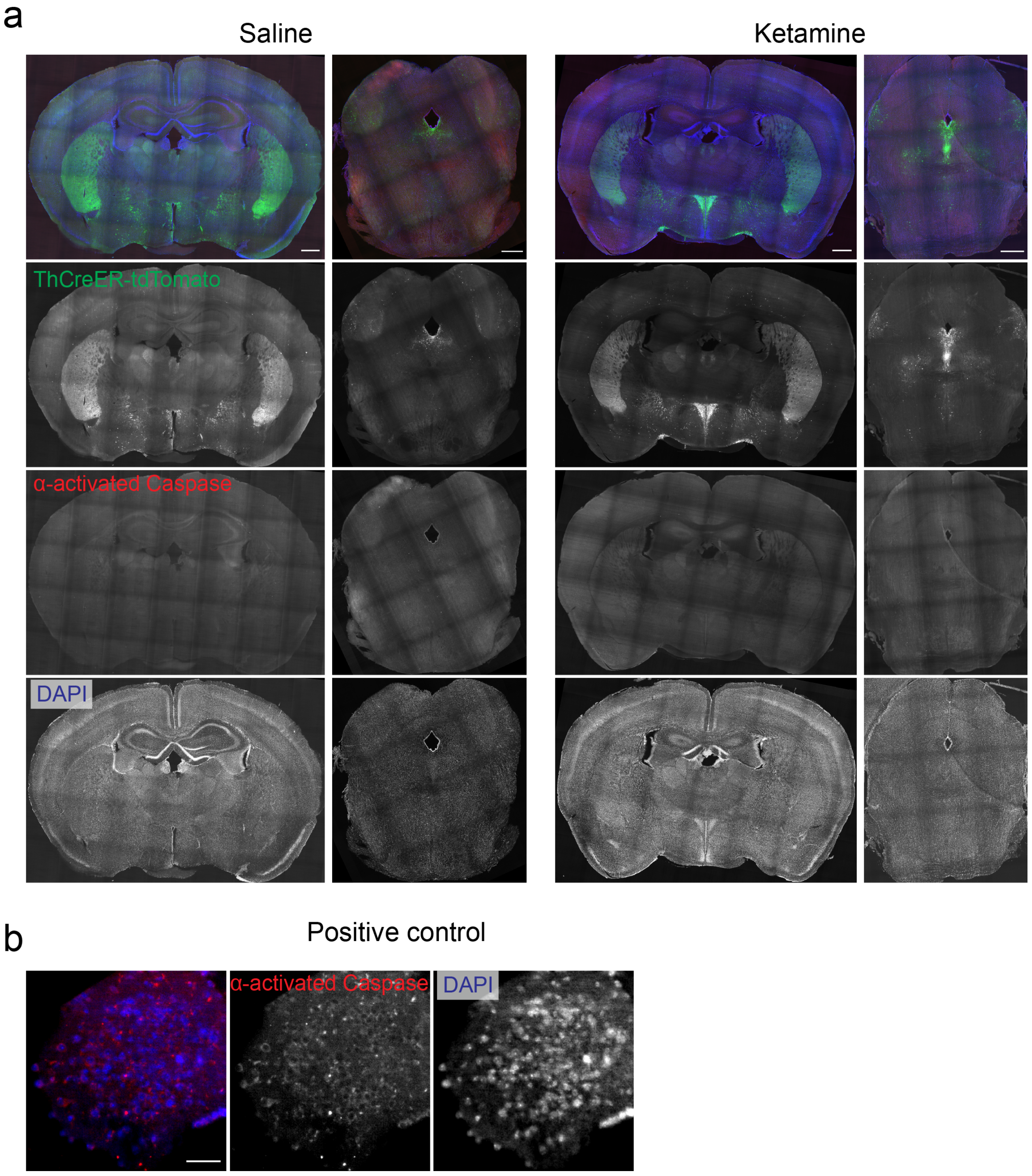


**Supplementary Figure 1. Activated caspase immunostaining.** (a) Representative images showing activated caspase immunostainings in 10 day ketamine i.p. injected animals. (b) Positive control for activated caspase immunostaining on a stressed neuronal culture.

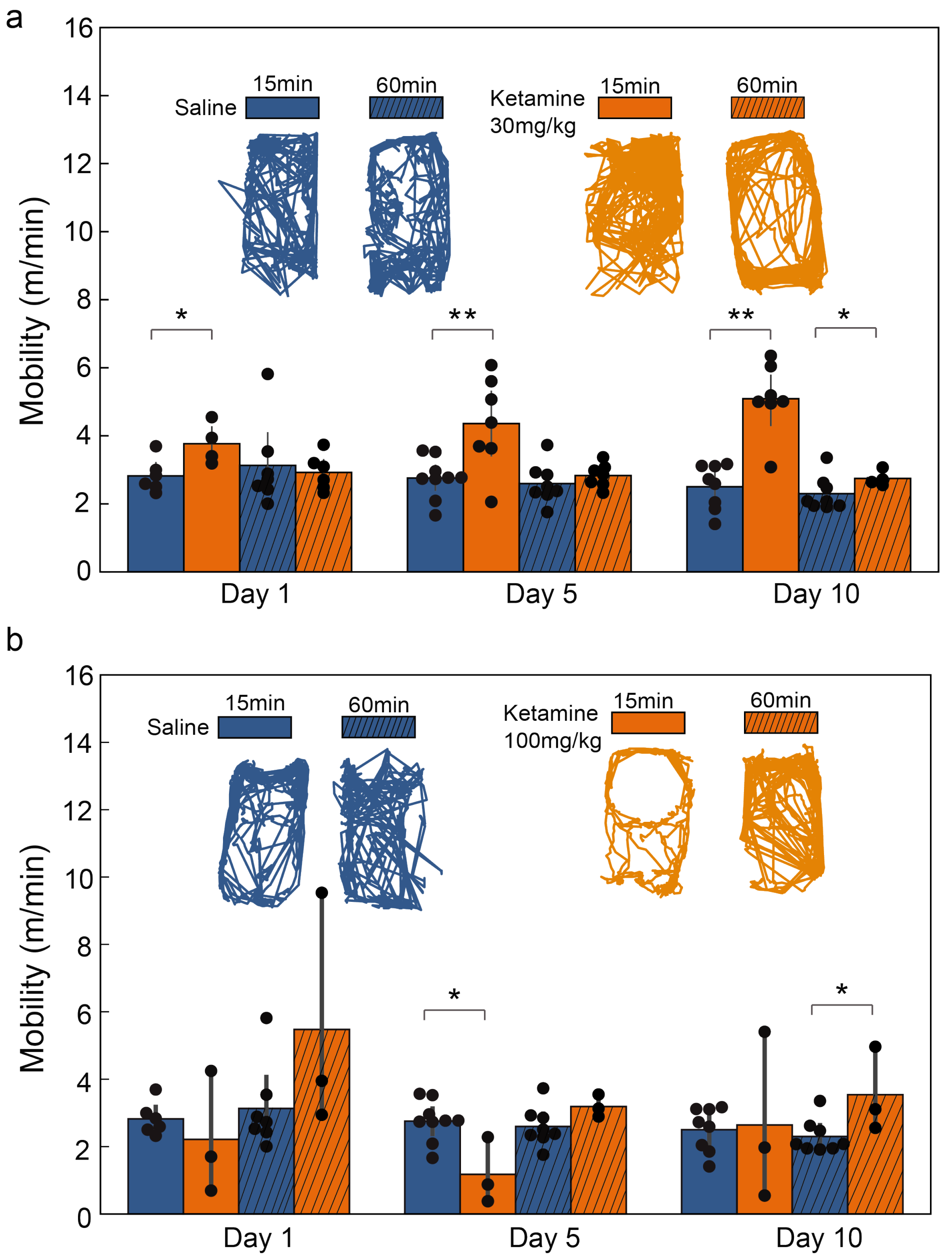


**Supplementary Figure 2. Locomotion behavior quantification**. (a) Quantification of mobility 15 and 60 minutes after daily i.p. injections of 30 mg/kg ketamine. (b) Quantification of mobility 15 and 60 minutes after daily i.p. injections of 100 mg/kg ketamine. Statistics was done using two-sided t-test. * denotes pvalue < 0.05, ** <0.01.

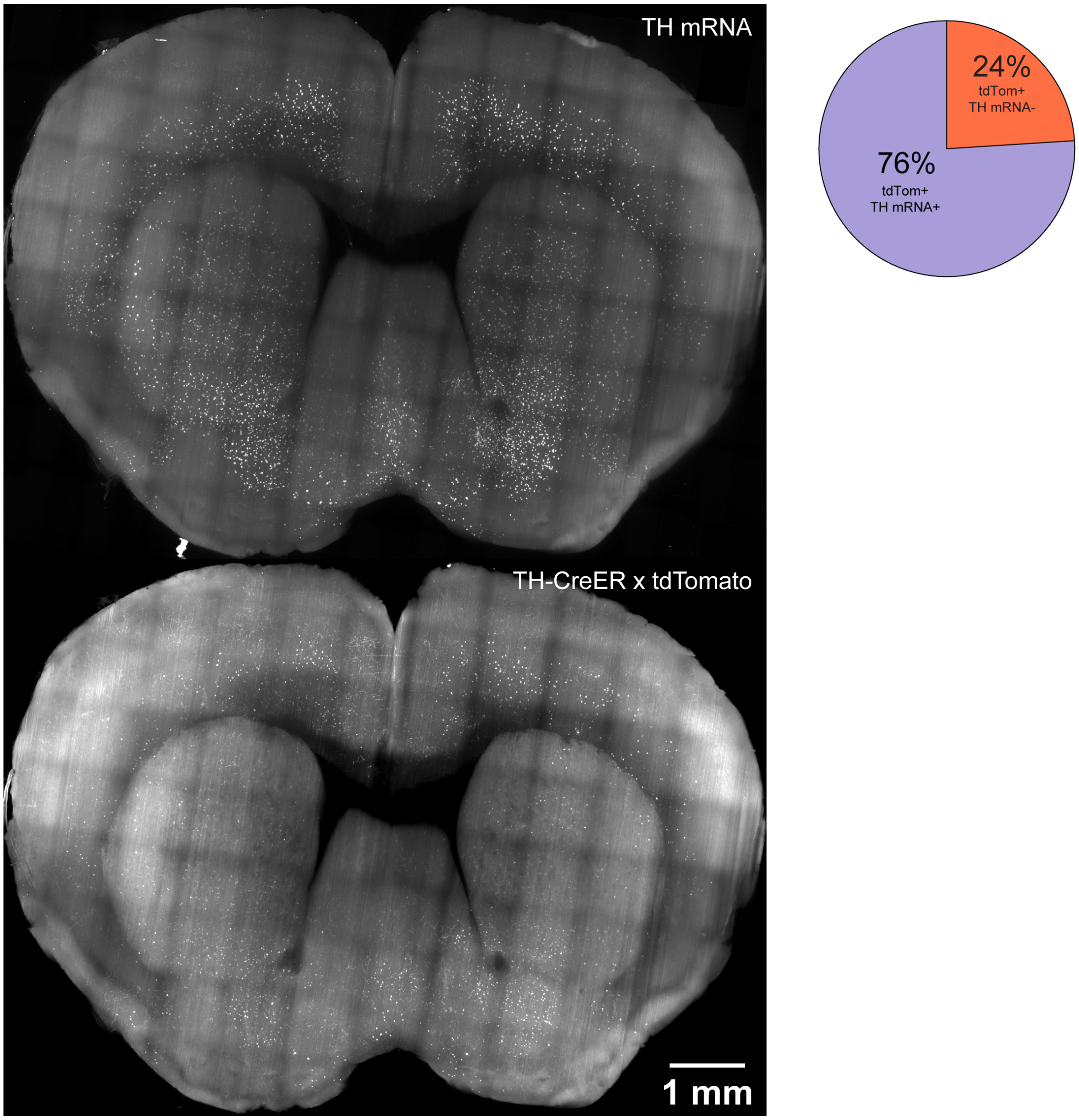


**Supplementary Figure 3. Characterization of TH-CreER transgenic line.** Co-localization of TH mRNA and TH-CreER x tdTomato signal. Pie chart quantifies the overlap of tdTomato signal with TH mRNA staining in all non-cortical regions.

**Supplementary Videos:**

**Supplementary Video 1. High-resolution whole brain TH immunostaining and imaging.** Representative example showing brain-wide TH+ neurons and their projections.

**Supplementary Video 2. Whole brain image registration.** Representative example of registration of a whole brain image to average reference brain.

**Supplementary Video 3. Volumetric rendering of whole brain TH+ cell segmentation.** Representative example showing brain-wide segmentation of TH+ neuronal cell bodies.

**Supplementary Video 4. Volumetric rendering of whole brain TH mRNA+ cell segmentation.** Representative example showing brain-wide segmentation of TH+ neuronal cell bodies.

**Supplementary Video 5. Comparison of whole brain TH mRNA+ (left) and TH protein+ neuronal cell bodies.** Representative examples comparing the distribution of TH mRNA+ and TH protein+ neuronal cell bodies.

**Supplementary Video 6. Whole brain heatmaps of alterations in TH+ neuron counts and projections.** Orange and cyan color-maps represent increases and decreases, respectively, after 10 days of ketamine (100 mg/kg) exposure.

**Supplementary Video 7. TH+ projection density changes in the prelimbic area.** Representative data of increases in TH+ projection densities in the prelimbic area after 10 days of ketamine (100 mg/kg) exposure. Cyan and magenta outline the prelimbic area and increased projection density region, respectively. Scale bar is 100 µm.

**Supplementary Video 8. TH+ projection density changes in the orbital area.** Representative data of increases in TH+ projection densities in the orbital area after 10 days of ketamine (100 mg/kg) exposure. Cyan and magenta outline the orbital area and the increased projection density region, respectively. Scale bar is 100 µm.

**Supplementary Video 9. TH+ projection density changes in the lateral amygdala.** Representative data of increases in TH+ projection densities in the lateral amygdala after 10 days of ketamine (100 mg/kg) exposure. Cyan and magenta outline the lateral amygdala and the increased projection density region, respectively. Scale bar is 100 µm.

**Supplementary Video 10. TH+ projection density changes in the dorsal auditory area.** Representative data of decreases in TH+ projection densities in the dorsal auditory cortex after 10 days of ketamine (100 mg/kg) exposure. Cyan and magenta outline the dorsal auditory area and the decreased projection density region, respectively. Scale bar is 100 μm.

**Supplementary Video 11. TH+ projection density changes in the postero-lateral visual area.** Representative data of decreases in TH+ projection densities in the postero-lateral visual area after 10 days of ketamine (100 mg/kg) exposure. Cyan and magenta outline the postero-lateral visual area and the increased projection density region, respectively. Scale bar is 100 μm.

**Supplementary Video 12. TH+ projections density changes in the Bed nuclei of the stria terminalis.** Representative data of decreases in TH+ projection densities in the Bed nuclei of the stria terminalis after 10 days of ketamine (100 mg/kg) exposure. Cyan and magenta outline the Bed nuclei of the stria terminalis and the increased projection density region, respectively. Scale bar is 100 μm.
